# Development of chemosynthetic microbial communities in organic falls deployed in the deep Southwest Atlantic Ocean

**DOI:** 10.1101/2021.02.16.431415

**Authors:** Augusto M. Amendola, Francielli V. Peres, Julio C. F. Moreira, Paulo Y. G. Sumida, Fabiana S. Paula, Vivian H. Pellizari

## Abstract

The assembly and successional processes of microbial communities inhabiting deep-sea whale and wood falls are highly complex and vastly unknown, as a myriad of factors may affect the development of a chemosynthetic-based ecosystem on these organic islands. The chemoautotrophy supported by organic substrates is the basis of long-lasting ecosystems, considered biodiversity hotspots in the oligotrophic deep sea. Understanding how these microbial communities develop and the factors affecting them could shed light on processes related to the maintenance of biodiversity in this environment. We performed a whale- and wood-fall experiment in the southwest Atlantic on the Brazilian continental margin and investigated biofilm-forming bacterial and archaeal communities colonising these substrates, deployed at 1500 and 3300 m depth. The composition of the prokaryotic communities shared some similarities with previously reported organic falls in the north Pacific and the Mediterranean Sea, mainly regarding sulphur oxidising chemolithotrophic taxa from the phyla Campylobacterota and Proteobacteria. Communities were found to be highly different between the organic substrates, as whale fall associated biofilms presented a higher dominance of sulphur oxidising chemolithotrophs. We also observed a significant difference between the two sites, with the whale associated communities at the 1500 isobath presenting a faster establishment of the chemosynthetic taxa.

## INTRODUCTION

Communities inhabiting the deep sea are mostly dependent on organic matter derived from the euphotic zone and often endure low carbon input conditions (Goffredi and Orphan 2010; Smith 2012). Large animal carcasses and wood logs can sink rapidly and, once landed in the deep sea, they provide substantial inputs of locally restricted organic substrate, thus promoting significant changes in the surrounding environment (Wolff 1979; Smith and Baco 2003). Whale and wood falls may persist for decades in the bottom of the ocean (Smith and Baco 2003; Palacios *et al.* 2009), feeding and sheltering vast microbial and faunal communities in complex ecological successional processes. With time, the decay of the organic matter leads to the build-up of reduced chemicals, which promote the development of chemosynthesis-based ecosystems (Jørgensen and Boetius 2007), with communities sharing similarities with those found in hydrothermal vents (Bernardino *et al.* 2010).

While in hydrothermal vents abiotic reactions are the source of reduced chemicals (Dick 2019), in organic falls these molecules are formed as a result of the microbial degradation of the organic matter. Small organic compounds released from the lipid rich bones (Bennett *et al.* 1994; Smith *et al.* 2014, 2015) or from the wood matrix (Laurent *et al.* 2013; Kalenitchenko *et al.* 2015) are anaerobically degraded by sulphate-reducing bacteria. This activity leads to the build-up of sulphide in the surrounding environment (also called “sulphophilic stage”), which promotes the establishment of sulphide-oxidising chemosynthetic microbes (Baco and Smith 2003; Schuller, Kadko and Smith 2004; Treude *et al.* 2009). This process of primary productivity supports the growth of heterotrophic microorganisms and specialised fauna(Bernardino *et al.* 2012). In wood falls, microbes capable to degrade the lignocellulosic substrate (Boer *et al.* 2005; Palacios *et al.* 2009; Bienhold *et al.* 2013; Fagervold *et al.* 2013; Ristova *et al.* 2017), and wood-boring bivalves harbouring symbiotic bacteria in their gills (Turner 1973; Voight 2009; Saeedi *et al.* 2019) seem to play an important role breaking down recalcitrant material releasing labile substrates.

Whale and wood falls have been coined as “organic islands” that contribute to the dispersal of macrofauna along different deep-sea chemosynthetic ecosystems (Smith *et al.* 1989; Distel *et al.* 2000; Sumida *et al.* 2016). Whale falls are estimated to occur in high frequency, especially in migratory routes (Smith *et al.* 1989; Butman, Carlton and Palumbi 1995; Perry, DeMaster and Silber 1999; Simabukuro *et al.* 2020). In addition, wood remnants are widely present in the marine environment, particularly on the continental margins, and may reach bathyal depths through underwater canyons and distant abyssal areas (Vetter and Dayton 1998; Amon *et al.* 2017). Hence, these long-standing organic islands not only provide a source for the enrichment of reduced substrates to fuel primary productivity in the oligotrophic ocean floor, but also function as important habitats for the biogeography of a diversity of organisms. Therefore, there is a demand for studies investigating the development of these chemosynthetic microbial communities on organic falls. Of particular interest is whether the geographic location has a significant impact on these processes. Sumida *et al.* (2016) reported similarities among faunal communities inhabiting whale falls in the Atlantic and Pacific oceans. Such comparison on the chemosynthetic microbial communities has not been performed to date, as most surveys are concentrated in the north Pacific Ocean and Mediterranean, and mainly focused on the macrofauna characterisation (e.g.: Baco and Smith 2003; Goffredi *et al.* 2004; Lundsten *et al.* 2010; McClain and Barry 2014), with lower efforts to describe microbial composition (Bienhold *et al.* 2013; Fagervold *et al.* 2013; Cavalett *et al.* 2017; Ristova *et al.* 2017; Taboada *et al.* 2020). To date, in the south Atlantic, only one study investigated the bacterial community in one sediment sample underneath a whale carcass (Cavalett *et al.* 2017). In addition to geographic location, local factors such as seabed depth and regional environmental characteristics may play major roles in the microbial assemblage, affecting community composition and succession rate. Even though a few studies have attempted to decipher these environmental features, the results are contrasting (Palacios *et al.* 2009).

Aiming to bridge this gap, we performed a whale and wood fall experiment in the southwest Atlantic, on the Brazilian continental margin. Experimental deployment of organic substrates on the deep sea have been shown to be an efficient approach to study the community assemblages in these ecosystems (Baco and Smith 2003; Fagervold *et al.* 2013; Souza *et al.* 2020). We describe, for the first time in the south Atlantic deep sea, biofilm-forming bacterial and archaeal communities colonising whale vertebrae and wood parcels, deployed at 1500 and 3300 m depth. In addition, we used a synthetic substrate as an inorganic control to verify the community assemblage in the absence of organic matter. We tested whether chemosynthetic microbial communities could be established in organic substrates after 23 months in the Atlantic deep sea. The time frame for the development of a chemosynthetic community in organic substrates in the deep-sea is still under discussion. Treude *et al.* (2009) estimated that in about seven years these organic islands can originate chemosynthetic environments similar to those found in hydrothermal vents (Treude *et al.* 2009; Laurent *et al.* 2013; Yücel *et al.* 2013). However, this process may vary considerably for different locations with distinct environmental conditions. We hypothesised that the assemblage of the microbial communities on the biofilms would be dependent on the substrate type and also distinct at the experimental sites established at different depths.

## MATERIAL AND METHODS

### Experiment design and sampling

This study was part of The BioSuOr (Biodiversity and Connectivity of Benthic Communities on Organic Substrates in the Deep SW Atlantic Ocean) project (Barroso *et al.* 2018), established in 2013 in the South-eastern Brazilian continental margin. The area comprises a variety of benthic habitats, including submarine canyons, cold-water coral reefs, and seeps (Sumida *et al.* 2004; Bernardino, Berenguer and Ribeiro-Ferreira 2016; Almada and Bernardino 2017; Bernardino and Sumida 2017; Fujikura *et al.* 2017). In the deep Brazilian margin, three major water masses have been described: Antarctic Intermediate Water (AAIW), North Atlantic Deep Water (NADW), and Antarctic Bottom Water (AABW) (Durrieu De Madron and Weatherly 1994; da Silveira *et al.* 2000).

Two free-fall landers were deployed at the 1.500 meters (21° 27 ′0.36″S, 39° 53′ 47.4″W) and 3.300 meters (−22° 50′ 27,117 ″S, −38° 24′ 58.7988″ W) isobaths in Bacia de Campos, containing six parcels of each substrate: WH - Humpback whale vertebrae (*Megaptera novaeangliae*); WO - wood parcels (*Pinus elliottii*, ~ 30 cm diameter and 10 cm high); and SY - synthetic material (vinyl sponge pool mat) as a control (**Figure 1**). The substrates were interspersed in the compartments (with a 500 μm mesh bag) and lids. The lids remained open during the experiment to keep the substrates exposed to the water flow. Further details on the experiment design are described by (Saeedi *et al.* 2019). The substrates remained submerged for 23 months (June 2013 – May 2015). Thereafter, the landers were rescued through acoustic realising. The biofilms found in the organic falls were collected with sterile spatulas (**Figure 1C-D**) and stored at −80° C. As there was no visible biofilm formation on the synthetic substrates (**Figure 1E**), small portions of the parcels were collected for analyses. From the 1,500 m isobath lander, we collected eight samples from the whale vertebrae, five from wood plots and three from synthetic parcels. From the 3,300 m isobath, we analysed three samples from the whale vertebrae, three from the wood plots and two from synthetic plots, adding up to 23 samples. Sample numbers and environmental characterisation of the sites are presented in **Table S1**.

**Figure 1.**
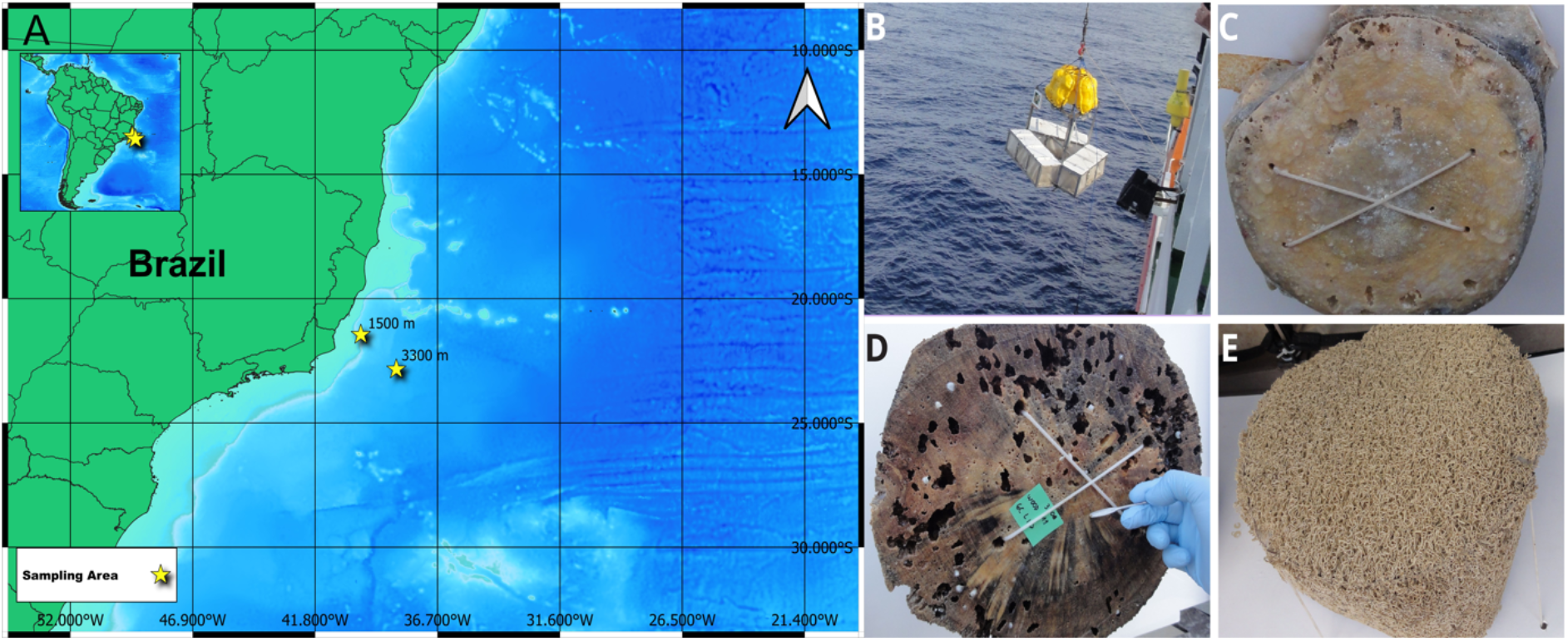
BioSuOr experiment. A) Map with the geographic location of the experimental areas at 1500 and 3300 m isobaths in the South-eastern Brazilian continental margin. The map was created using the QGIS software (QGIS Development Team, 2015-QGIS Geographic Information System. Open Source Geospatial Foundation Project. http://qgis.osgeo.org), with bathymetric data from ETOPO1 (https://www.ngdc.noaa.gov/mgg/global/) and world borders from (http://thematicmapping.org/downloads/world_borders.php). B) Deployment of the lander system consisting of a triangular aluminium frame containing a set of whale vertebrae, wooden block and synthetic carpet packed in mesh bags. C-E) Substrates parcels collected from the landers after 23 months submerged: C, Humpback whale vertebrae (*Megaptera novaeangliae*); D, wood (*Pinus elliottii*); E, synthetic (vinyl sponge pool mat).

### DNA extraction and 16S rRNA gene sequencing

Genomic DNA was extracted from 0.2 g (wet weight) of biofilm (or equivalent weight of pieces of the synthetic material) using the Power Biofilm DNA Isolation Kit (MoBio Laboratories, CA, USA). DNA concentration was assessed with Qubit dsDNA HS assay kit (Thermo Fisher Scientific, São Paulo, Brazil), following the manufacturer’s instructions, and measured using a Qubit Fluorometer 1.0. DNA integrity was verified in 1% agarose gel. The hypervariable regions V3 and V4 of the ribosomal 16S gene were amplified using the universal primers 515F (5’-GTGYCAGCMGCCGCGGTAA-3’) and 926R (5’-CCGYCAATTYMTTTRAGTTT-3’). The PCR reaction started with denaturation of 95 °C for 3 minutes, followed by 35 cycles at 95 °C for 30 seconds, annealing at 57 °C for 30 seconds, extension at 72 °C for 30 seconds and final extension at 72 °C for 5 minutes. A second PCR reaction was performed to add indices to the amplicons, followed by purification, quantification and normalisation of the libraries. The libraries were sequenced in the Illumina Miseq platform (2 x 250pb) at Mr DNA/Molecular Research (Shallowater, TX, USA), following standardised procedures. Data Availability of the datasets generated during the current study are available in the GenBank repository under BioProject ID PRJNA675849.

### Data analysis

Processing and quality control of 16S rRNA reads was performed using the QIIME2 platform version 2019.4 (Bolyen *et al.* 2019). After demultiplexing, the quality of the reads was checked and processed using the DADA2 algorithm (Callahan, McMurdie and Holmes 2017) implemented in a QIIME2 plugin. Forward and reverse raw sequences were trimmed, filtered and denoised to exclude poor quality reads. The reads were then merged, and the chimera were removed. The Amplicon Sequence Variants (ASVs) were generated following default parameters and classified using Silva v.138 taxonomy database (Klindworth *et al.* 2013).

Prior to calculating the alpha and beta diversity the ASV table was normalized by variance stabilising transformation using the package *DESeq2* (Love, Huber and Anders 2014a). Alpha diversity indexes (Chao1 and Shannon) were calculated using the package *phyloseq* version 1.28.0 (McMurdie and Holmes 2013). Two-way ANOVA was used to test the effects of depth and substrate type on the indexes, using the package *agricolae* version 1.3-1 (Mendiburu and Yaseen 2020). Non-Metric Multidimensional Scaling (nMDS) analysis with Bray-curtis as dissimilarity index was carried out using *vegan* package version 2.5-6 (Oksanen et. al., 2018). PERMANOVA analysis(Anderson 2001) was performed to investigate the differences in communities across substrates and depths. To identify the ASVs significantly more abundant across substrate types and depths, we performed *DESeq2* - Differential Expression analysis implemented for community sequence count data (Love, Huber and Anders 2014b). Core Microbiome analysis within and across substrates was carried out using the *lyr* package (Wickham 2011).The ASVs shared by all substrates were considered the core microbiome. The functional potential of the communities was predicted using the Functional Annotation of Prokaryotic Taxa (FAPROTAX) database version 1.2.4 (Louca, Parfrey and Doebeli 2016). All statistical analyses and plots were performed using R version 3.6.1 (R Core Team, 2019).

## RESULTS

### Taxonomic profiling of the prokaryotic communities

The composition of the prokaryotic communities varied widely across samples collected from the different substrates. Non-metric multidimensional scaling analysis on the community composition indicated that the samples tended to cluster by substrate and by depth (**Figure 2**). We observed that whale bone-associated biofilms grouped distantly from the other substrates. According to PERMANOVA analysis 21% of the community variability was explained by the substrate type (p<0.001), and less than 7% by depth (p<0.05), while the interaction substrate x depth accounted for 15% of the variability (p<0.05) (**Table 1**).

**Figure 2.**
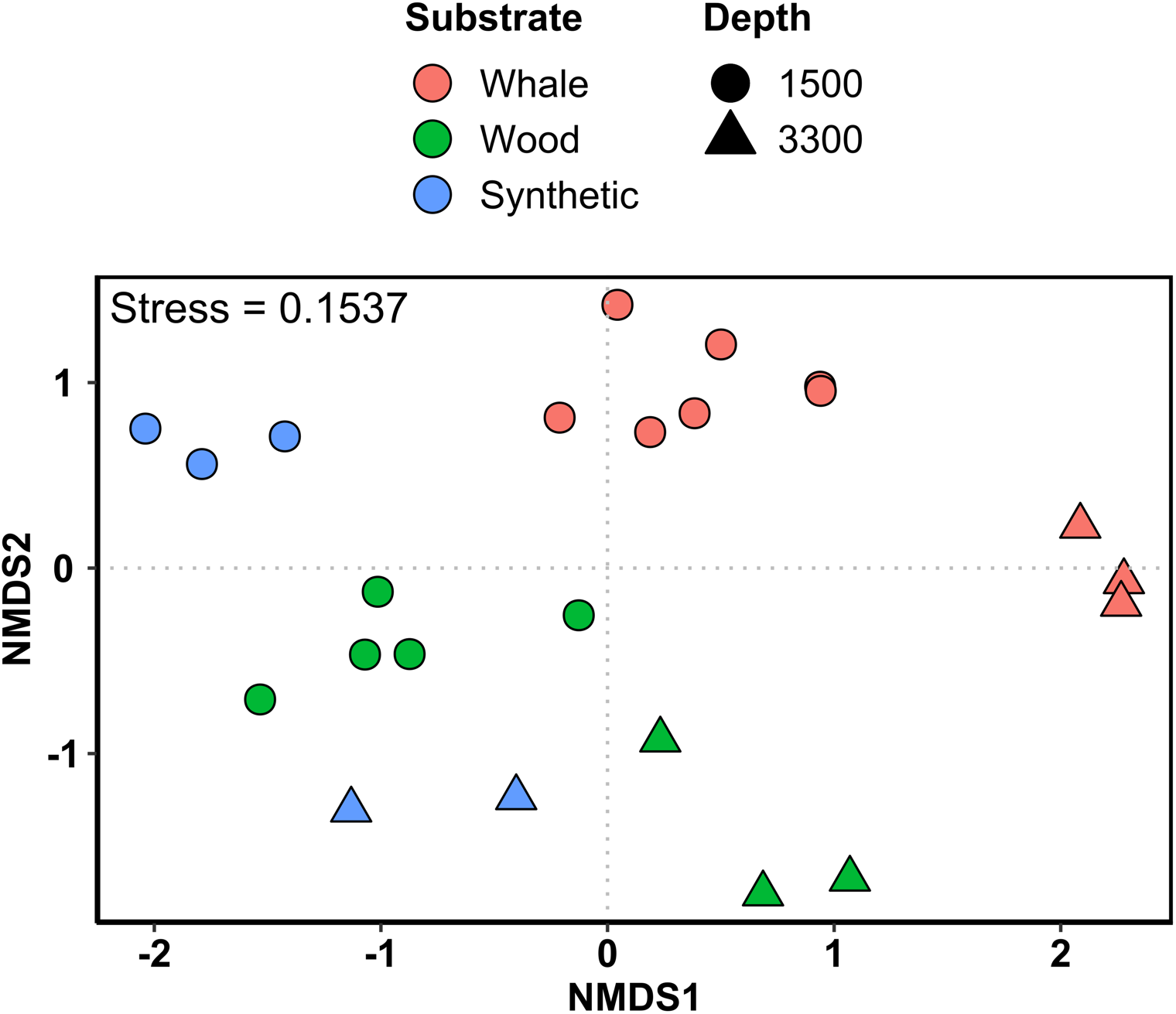
Non-metric multidimensional scaling of the prokaryotic communities in the samples from the different substrates. Distance index: Bray-Curtis.

**Table 1.** PERMANOVA analysis of the microbial communities at different substrates and depth

| Factor | F | R<sup>2</sup> | P |
| --- | --- | --- | --- |
| Substrate | 2.092 | 0.210 | **0.001** |
| Depth | 2.037 | 0.068 | **0.002** |
| Substrate * Depth | 1.527 | 0.153 | **0.003** |

All communities were largely dominated by Bacteria. The proportions of reads assigned to the Domain Archaea were below 0.6% and 1.4% in samples from 1500 and 3300 m, respectively. Most archaeal sequences were assigned to the phylum Nanoarchaeota (**Figure 3**), comprising all archaeal reads in several samples from different substrates. The phylum Crenarchaeota was also detected in some samples, mainly from the synthetic substrate (**Figure S1**).

**Figure 3.**
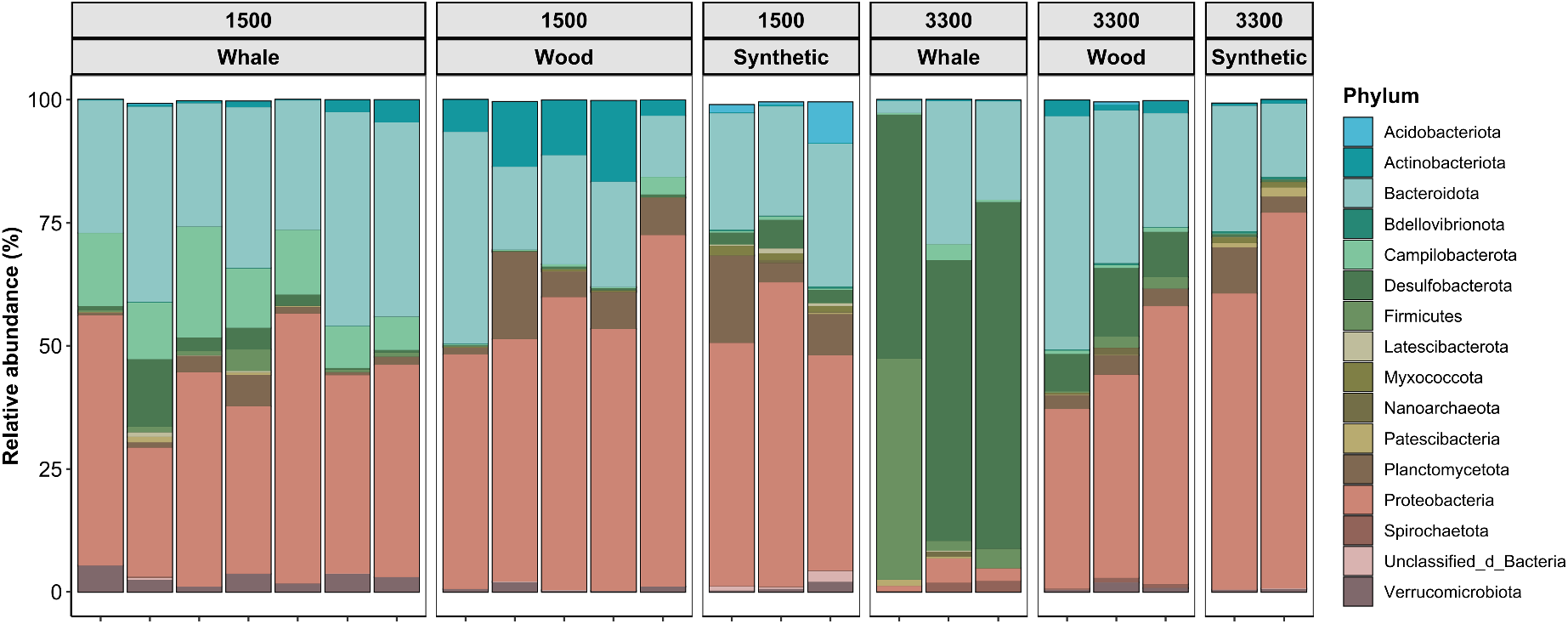
Relative abundance of archaeal and bacterial phyla in samples collected from the different substrates at the 1500 and 3300 m isobaths. Phyla with relative abundance above 0.1% are included in this figure. Rare groups are presented in Figure S1.

In the microbial communities from the 1500 m isobath, Proteobacteria and Bacteroidota were found to be dominant groups in all substrates, with relative abundance values above 48 and 28%, respectively. ASVs assigned to Planctomycetota were also abundant in most samples, mainly in synthetic (SY) and wood (WO) substrate communities, reaching up to 10 and 9% in those substrates, respectively. In whale vertebra (WH) biofilms, we observed high proportions of the phylum Campilobacterota (up to 22%). By contrast, Actinobacterota represented up to 10% of the communities in the WO biofilms, while it was observed in very low abundance in the other substrates **(Figure 3A)** - this was confirmed by analysis of differential abundance, using *DESeq2*, which showed that ASVs assigned to this phylum were significantly in higher proportions in WO samples (**Figures S2B-C**). In the SY communities, Acidobacteriota accounted for up to 8%, while this phylum was below 0.2% in the other substrates. Finally, the phylum Desulfobacterota was detected in all samples, and contributed to up to 13% of the communities (in WH) from the 1500 m site (**Figure 3; Figures S2A-B**).

The WH communities from the 3300 m isobath presented the most remarkable differences, when compared to the other substrates - they were also notably distinct from WH at 1500 m. The 3300 m WH communities were largely dominated by the phylum Desulfobacterota (50 to 70%), with Bacteroidota and Firmicutes also playing expressive contributions. Proteobacteria was poorly represented in these WH samples, while it was among the dominant phyla in WO e SY substrates from the same depth, along with Bacteroidota, Desulfobacterota and Planctomycetota. As observed for the 1500 m samples, Actinobacteria was found to be more associated with the 3300 m wood biofilms, despite lower relative abundance at this depth (**Figure 3**).

Similar to the patterns observed for community composition, substrate type was the strongest factor affecting richness and diversity, as assessed by Chao1 and Shannon indexes, respectively (**Figure S5; Table S2**). The factor depth alone did not cause significant change in either richness or diversity. Overall, the communities found in the synthetic substrate, mainly from 1500 m depth, tended to present the highest indexes, while the lowest values were observed for whale bone biofilm from the 3300 m isobath.

### Predicted functions

The 4172 ASVs detected across the samples were searched against the FAPROTAX database (Louca, Parfrey and Doebeli 2016) to explore their functional potential based on phylogenetic relations with cultured representatives. The predicted metabolisms varied widely throughout the samples and included functions related to sulphur and nitrogen cycling, carbon utilisation and energy production, as well as interactions with other organisms **(Figure 4)**. Sulphur compounds respiration and oxidation were predicted for all substrates, but these functions were notably at higher proportions for the whale bone associated biofilms. It was noteworthy that processes related to sulphur oxidation were more associated with 1500 m WH biofilms, while S reduction was associated with the 3300 m WH communities. Nitrate reduction and fermentation were predicted across all samples, suggesting the importance of other anaerobic metabolisms. Methanotrophy was mainly associated with 3300 m WH samples, while other one-carbon metabolisms were predicted for the wood and synthetic substrates.

**Figure 4.**
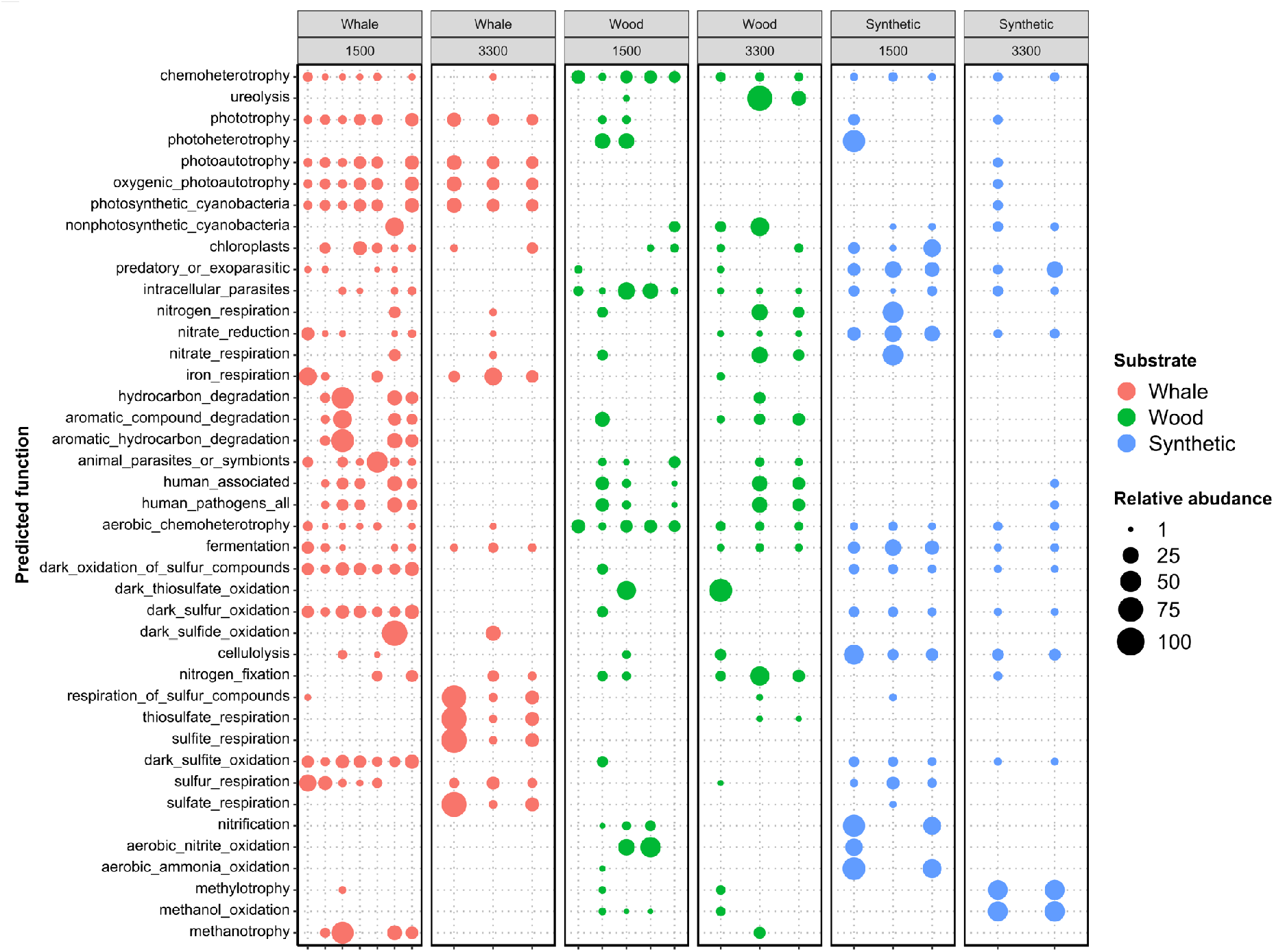
Predictions on the potential metabolic capabilities harboured by the microbial communities associated to each substrate at 1500 and 3300 m depth. The functions were predicted by FAPROTAX analysis. The data was normalised by function. The size of the circles indicates the relative abundance of each function across samples. Colours refer to substrate type.

We further analysed the relative abundance of microbial taxa with reported capability to perform reduction and oxidation of sulphur compounds. We found 272 ASVs assigned to 23 families known to comprise organisms that carry the potential for such metabolisms (**Figure 5**). In agreement with the functional predictions inferred by FAPROTAX analysis, WH-associated biofilms presented the highest relative abundance of those groups. Bacterial families containing sulphate reducing (SR) taxa were found to be among the dominant groups in several organic samples, mainly those from 3300 m, where it accounted for up to 70% of the WH communities (**Figure 5A**). At 1500 m, WH-associated SR bacteria were mostly composed by the family Desulfocapsaceae, followed by Geopsychrobacteraceae. In addition, at 1500 m, most ASVs assigned to these families were present in significantly higher proportions in WH (**Figure S2A**). Surprisingly, low proportions of SR bacteria were observed in 1500 m WO, with values even lower than what was observed for SY. At the 3300 m isobath, Desulfocapsaceae was the most abundant SR family across all organic substrates (with up to 60% of overall relative abundance in WH), followed by Desulfobacteraceae and Geopsychrobacteraceae (these two families were mainly observed in whale bone associated biofilms). *DESeq2* analysis indicated significantly higher proportions of SR ASVs in the WH and WO communities, in relation to SY (**Figure S3A-B**).

**Figure 5.**
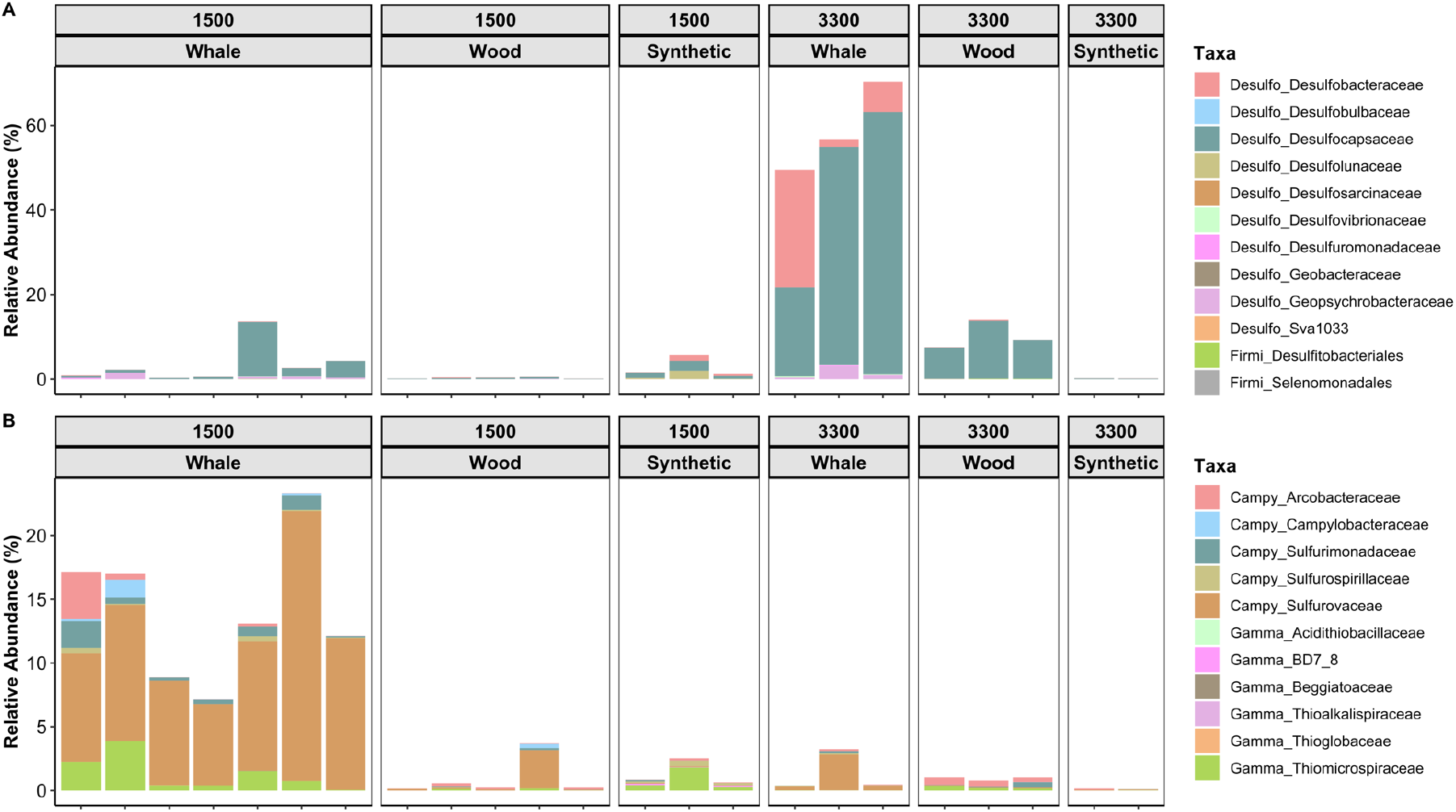
Relative abundance of bacterial taxa with reported function in S compounds reduction (A) and oxidation (B) in samples collected from the different substrates at depths of 1500 and 3300 meters.

Regarding the chemolithoautotrophic sulphur oxidising groups, differently from SR, we observed the highest relative abundances in samples from 1500 m, mainly in WH-associated biofilms (**Figure 5B**). Most sulphur oxidising communities were dominated by the phylum Campylobacterota, which comprised five families. The family Sulforovaceae was detected in all substrates and in WH biofilms from 3300 m it represented between 10 and 20% of the overall prokaryotic communities. Other families from the phylum Campylobacterota were found to be present in high relative abundance across the samples, including Arcobacteraceae and Sulfurimonadaceae. *DESeq2* analysis indicated significantly higher proportions of most ASVs assigned to these taxa in WH substrate (**Figure S2 and S3**). We also detected six Gammaproteobacteria families containing groups with reported capability to perform chemolithoautotrophic sulphur oxidation (**Figure 5B**). These included Thiomicrospiraceae, known to comprise endosymbiotic species, which was widespread among all substrates at 1500 m, while at 3300 m it was mainly associated with wood biofilms.

### Core Microbiome

The core microbiome analysis was used to identify ASVs shared by all samples within and between substrate types, regardless depth. Due to the different number of samples analysed for each substrate type, direct comparisons on the sizes of the core microbiomes may not provide meaningful information. Hence, only qualitative comparisons will be presented.

Two ASVs were found to be shared among 100% of the samples analysed. They were assigned to the genera *Sulfurovum* (class Campylobacteria) and *Sedimentitalea* (class Alphaproteobacteria). In addition, ASVs assigned to the following taxa were identified as core microbiomes between pairs of substrates: WH/WO - *Desulforhopalus* (Desulfobulbia) and Sphingomonadaceae (Alphaproteobacteria); WH/SY - *Clostridium* (Clostridia) and Thiotrichaceae (Gammaproteobacteria); and WO/SY - *Pseudovibrio* and *Sedimentitalea* (both from class Alphaproteobacteria).

Among the 12 ASVs shared by all whale bone biofilm communities, we observed several groups with reported capability for S reduction (genera *Desulfobacter* and *Desulfuromusa*, and family Desulfocapsaceae) or oxidation (genus *Sulfurovum*). By contrast, among the 30 ASVs that composed the core microbiome of the samples from the synthetic substrate, only one was related to the sulphur metabolism (genus *Sulfitobacter*). Interestingly, only three ASVs were exclusive to the WO core microbiome, and none of them were related to the S metabolism (**Table 2**).

**Table 2.**
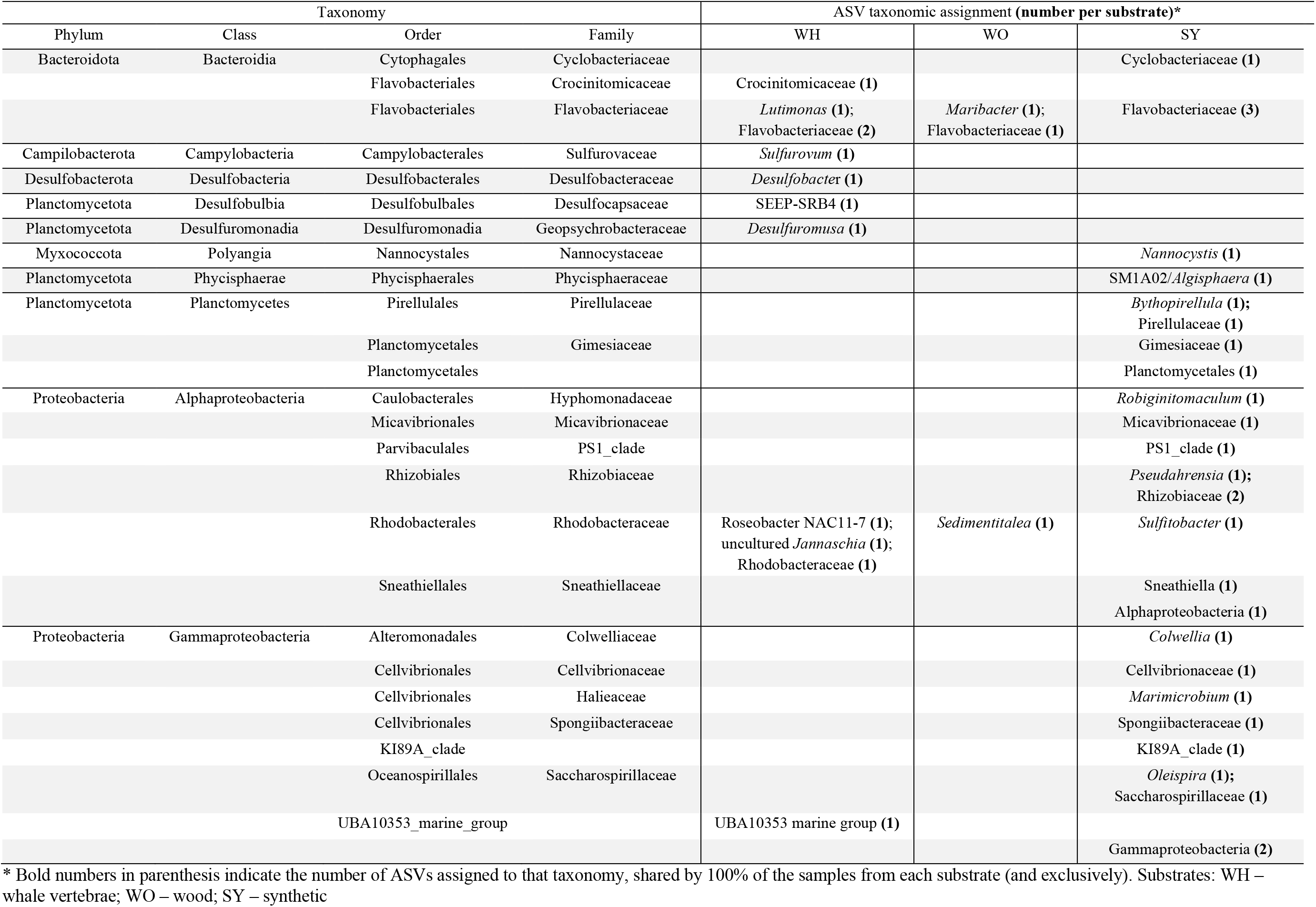
Core Microbiome analysis – list of ASVs shared by (and exclusive to) all samples from each substrate.

## DISCUSSION

We evaluated the potential for the development of chemosynthetic microbial communities on organic falls experimentally deployed in deep-sea areas, in the Southwest Atlantic. A diverse fauna associated with these organic falls was also retrieved, and it has been described elsewhere (Barroso *et al.* 2018; Saeedi *et al.* 2019; Souza *et al.* 2020). High throughput sequencing of the 16S rRNA gene indicated a high diversity of prokaryotes in the samples spanning over 40 phyla. Substrate type was found to be a significant factor driving changes in community diversity and composition in the biofilms collected after 23 months from the deep-sea. Organic matter type is known to affect microbial community assembly (Fagervold *et al.* 2014). In addition, as the decomposition process proceeds, microbial diversity is expected to alter in response to changes in substrate availability and in the characteristics of the surrounding environment (Kalenitchenko *et al.* 2016). These shifts may favour specialised groups, which can thrive and become dominant taxa. In fact, in the 1500 m samples, WH and WO communities tended to have lower microbial richness and diversity, in comparison to the control SY. WH and WO communities were enriched with organisms related to organic matter decay, as well as chemolithoautotrophic groups (from Campilobacterota and Proteobacteria phyla) (Borchert *et al.* 2020). In addition, there was an increase in Actinobacterota in WO, known for the capabilities to degrade recalcitrant organic matter (Bull *et al.* 2005; Steger *et al.* 2007). Interestingly, at the 3300 m depth, this decline in diversity was only observed for WH samples. A very low diversity in those communities was observed, which is likely to be an effect of the expressive dominance of few groups, primarily from the phylum Desulfobacterota, which comprises a diversity of sulphate reducing bacteria (Waite *et al.* 2020).

The composition of the microbial biofilms formed on the organic substrates, mainly in WH samples, was comparable with those found in chemosynthesis-based ecosystems. The lipid rich whale bones provide large amounts of labile substrates for microbial growth. High microbial activity can deplete the oxygen in the surrounding environment, favouring anaerobic metabolisms, such as fermentation, sulphate reduction and methanogenesis (Goffredi *et al.* 2008; Treude *et al.* 2009). Even in oxygenated waters, anoxic environments can occur in the proximity of the organic falls (Fagervold *et al.* 2012, 2013; Yücel *et al.* 2013; Ristova *et al.* 2017). Sulphate-reducing bacteria were found to be abundant and widespread in WH associated substrates. Their metabolism leads to the release of sulphide in the environment, which can be used as an energy source by the chemolithoautotrophic sulphide-oxidising microbes (Smith and Baco 2003; Treude *et al.* 2009; Bienhold *et al.* 2013). We detected a diversity of S oxidising bacteria distributed in 11 families from the taxa Campilobacterota and Gammaproteobacteria. *Sulfurovum* (Sulfurovaceae) was the most abundant genus and part of the core microbiome. This genus was among the dominant taxa in WH communities from 1500 m, and significantly more abundant in this substrate (**Figures S2 and S3**). Other Campylobacerota genera were also prevalent across the samples, including *Sulfurospirillum* (Sulfurospirillaceae) and *Sulfurimonas* (Sulfurimonadaceae). These three genera are often reported among the dominant groups in whale fall carcasses during the sulphophilic stage (Allison *et al.* 1991; Deming *et al.* 1997; Cavalett *et al.* 2017), in hydrothermal vents (He *et al.* 2015; Fortunato and Huber 2016; Meier *et al.* 2017; Cerqueira *et al.* 2018), and other sulphur rich environments (Yamamoto *et al.* 2010; Akerman, Butterfield and Huber 2013). Campylobacterota was also found to be a dominant group in the only microbiological survey on whale fall performed in the south Atlantic Ocean to date (Cavalett *et al.* 2017). The authors reported that sediment samples collected underneath a whale carcass found at 4200 m contained high proportions of Epsilonproteobacteria - the taxa previously assigned to this class are now included in the phylum Campylobacterota, according Silva v. 138 (Bizic *et al.* 2020). We also detected Gammaproteobacteria S-oxidisers, mainly from the family Thiomicrospiraceae. This family has been reported in hydrothermal vent communities (Li *et al.* 2020) and comprises groups that live in symbiosis with vent-associated fauna (Tourova *et al.* 2006).

The presence of recalcitrant molecules in wood logs, such as cellulose, hemicellulose and lignin (Florencio, Couri and Farinas 2012), may be associated with a slower decomposition of this substrate and consequent delay in the establishment of chemosynthetic communities, in relation to the WH communities. To hydrolyse these complex molecules, the activity of specialised organisms is required (Ljungdahl and Eriksson 1985) Cellulolytic activity has been reported in several microbial taxa, including Actinobacteria, Bacteroidota, Firmicutes, Proteobacteria and Spirochaetes (Carere *et al.* 2008). By contrast, lignin degradation is mostly restricted to fungi, and the known bacterial groups with this activity are mainly in the taxa Actinomycetes, α-Proteobacteria and γ-Proteobacteria (Bugg *et al.* 2011; Huang *et al.* 2013). We observed the above-mentioned bacterial taxa in the biofilms analysed, but only the Actinobacteria was found to be enriched in WO. However, as discussed by Fagervold *et al.* (2012, 2013) and Ristova *et al.* (2017), microbial communities in sunken wood parcels may be highly dynamic in relation to time and space. Therefore, different groups may contribute to higher proportions of the community in different phases of the decomposition process. Yet, the prevalence of Actinobacteria in the wood biofilms, mainly in samples from 1500 m may suggest the importance of this taxon for the degradation of recalcitrant molecules in the wood parcels. In addition. Saeedi *et al.* (2019) reported the presence of wood boring bivalves from the genus *Xylophaga* and the formation of burrows were observed in the wood parcels, indicating that the decomposition process was in progress. But as observed by Kalenitchenko *et al.* (2018b, 2018a), the production of sulphide in wood falls may be initiated before the colonisation by the bivalves.

The microbial community composition also changed significantly between the two experimental sites, at the 1500 and 3300 m isobaths. Local environmental characteristics are known to affect microbial diversity (DeLong 2006). In fact, we registered distinct values of temperature, pressure and DO between both experimental sites, which may have affected the communities. In addition, the two sites were under the influence of distinct water masses, North Atlantic Deep Water (NADW) and Antarctic Bottom Water (AABW) (1500 and 3300 m sites, respectively) (Durrieu De Madron and Weatherly 1994; da Silveira *et al.* 2000), which may have different abiotic and biotic composition - including microbial pools functioning as sources for the development of the new communities (Hamdan *et al.* 2013; Walsh *et al.* 2016). In relation to S cycling microorganisms, there was a noteworthy site-related shift in WH communities, as a higher enrichment of S oxidising groups was observed at 1500 m (**Figures 5B and S4A**), while S reducing bacteria were thriving at 3300 m (**Figures 5A and S4A**). The profile suggests that the establishment of the chemosynthetic communities at the site of the deeper isobath may have lagged behind the process in the shallower area. This delay in community succession could be related to a multitude of factors, including decreasing microbial biomass with geographic distance from shore (Kallmeyer *et al.* 2012). In wood associated communities, an enrichment of SRB was also observed at 3300 m, in relation to the control (SY). However, no increase in the S oxidising taxa was observed at any depth in WO. Longer experimental design with periodic sampling would be required to unveil the successional processes and environmental constraints at different depths. Yet, logistic complexities and high costs humper the conduction of similar experiments in deep-sea areas.

The time required for the establishment of a chemosynthetic community in organic substrates in deep-sea areas seems to be dependent on a series of factors. Testing different locations, depths and time, Fagervold *et al.* (2013) observed that the immersion period length was the most important factor affecting the microbial community composition in wood parcels. In experimental whale fall studies in Santa Catalina Basin, in the north Pacific, the formation of the sulphophilic stage was observed after 2 years (Baco-Taylor 2002; Smith, Baco and Glover 2002; Smith and Baco 2003; Bernardino *et al.* 2010). Here we reported the enrichment of related microbial communities in whale bones after 23 months in the south Atlantic and an effect of the ocean depth on this system. In coastal shallow sediments in the Antarctic ocean, a rapid development of chemosynthetic communities has been reported in whale bones experimentally sunk for three months at 20 meters, and for one year at 10 meters depth (Taboada *et al.* 2020). The authors observed the prevalence of Epsilonproteobacteria (Campylobacterota) and Desulfuromonadales in the microbial mats. Regarding wood substrates, Yücel *et al.* (2013) monitored a wood fall experiment for up to 330 days and reported late production of reduced S compounds. In a long-term monitoring experiment (up to 3 years), Ristova *et al.* (2017) described the successional assembly of the microbial communities in wood parcels sunk at 1700 m. The communities were initially dominated by Alphaproteobacteria, which were later outnumbered by Deltaproteobacteria and Epsilonproteobacteria. Other studies reported the dominance of Bacteroidota and proteobacteria in sunken wood parcels in the Mediterranean Sea and in the Pacific Ocean during earlier stages of decomposition (Fagervold *et al.* 2012, 2013). We also observed high proportions of Bacteroidota in the biofilms, not only in wood, but also in whale substrates, similarly to previous reports (Taboada *et al.* 2020). However, we could not relate the relative abundance of this clade to the successional stage of the communities.

In this work we report that organic falls-associated chemolithoautotrophic communities developed in the south Atlantic deep ocean share similarities with those reported for distant geographic locations, such as the north Pacific and the Mediterranean Sea. Yet, particularities of these communities in comparison to other areas highlight the complexity of the environmental constraints underpinning this process. The main factor affecting microbial community assembly was the type of organic matter, with a higher dominance of S-oxidising chemolithotrophs in whale associated biofilms. In addition, regional environmental characteristics of sites located at different depths seem to play a significant effect on the community successional process. The study sheds light on the microbial components of these organic islands in the south Atlantic deep sea and the formation of these important ecosystems of the oligotrophic deep sea. In addition, it allows to explore specialised microbial pools, which are otherwise present in very low abundance and may not be detected in marine surveys.

## Supporting information

Supplementary data Atlantic Organic Falls

## SUPPLEMENTARY DATA

This manuscript contains supplementary data.

## ACKNOWLEDGEMENTS

We are thankful to the research teams of Laboratótio de Ecologia de Microorganismos (LECOM-IO/USP), Laboratório de Ecologia e Evolução de Mar Profundo (LAMP-IO/USP), Rosa Gamba and Cristina R. Nakayama (UNIFESP) for their sampling and scientific support, and to the captain and the crew of R/V Alpha-Crucis (IO-USP, FAPESP Process number 2010/06147-5), R/V Alpha Delphini and research polar vessel Almirante Maximiano (H-41) for the deployment and the recovering of the landers.

