## Supplementary data Atlantic Organic Falls for "Development of chemosynthetic microbial communities in organic falls deployed in the deep Southwest Atlantic Ocean"

**Figure / Table** **Page**

Figure S1……………………………………………………2
Figure S2……………………………………………………3
Figure S3…………………………………………………...…4
Figure S4…………………………………………………...…5
Figure S5…………………………………………………...…6
Table S1………………………………………………...…..7
Table S2……………………………………………...……..8

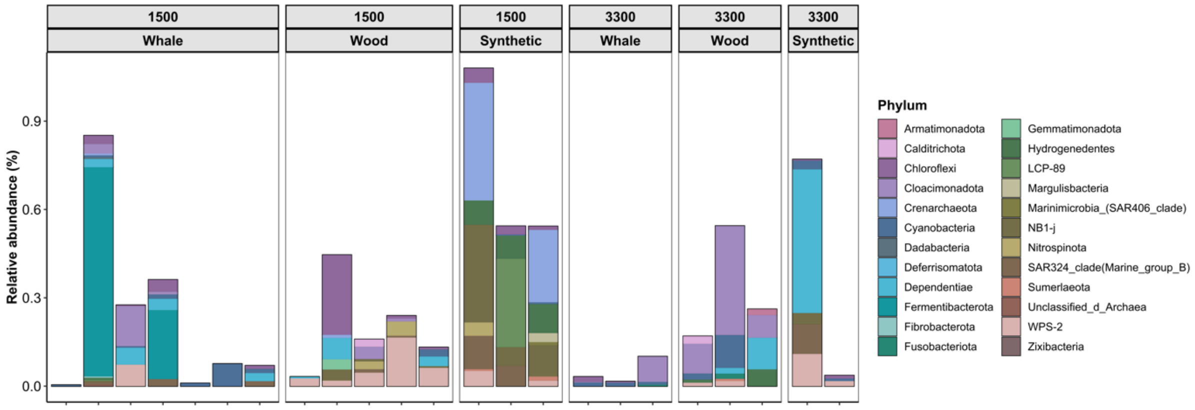


**Figure S1**. Relative abundance of rare bacterial and archaeal phyla in samples collected from the different substrates at the 1500 and 3300 m isobaths.

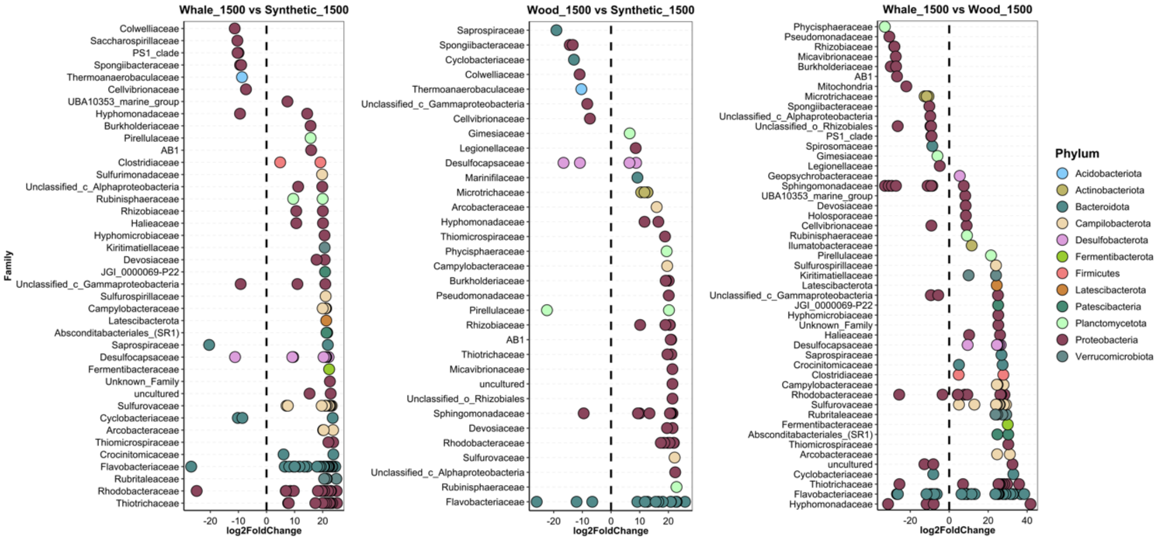


**Figure S2**. Differential abundance analysis of the ASVs between the substrate types at the 1500 m isobath. Log2FoldChange < 0 and > 0 indicates, respectively, ASVs with significantly lower and higher abundance in substrate 1 vs substrate 2: A) whale vs synthetic; B) wood vs synthetic; C) whale vs wood. The ASVs are grouped by family and coloured by phylum.

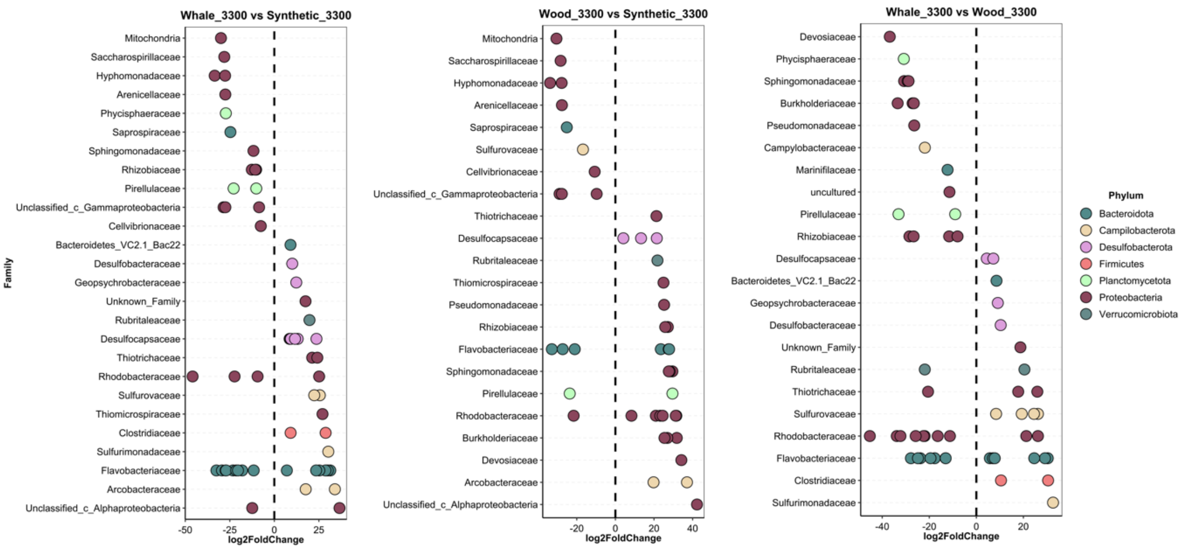


**Figure S3**. Differential abundance analysis of the ASVs between the substrate types at the 3300 m isobath. Log2FoldChange < 0 and > 0 indicates, respectively, ASVs with significantly lower and higher abundance in substrate 1 vs substrate 2:  A) whale vs synthetic; B) wood vs synthetic; C) whale vs wood. The ASVs are grouped by family and coloured by phylum.

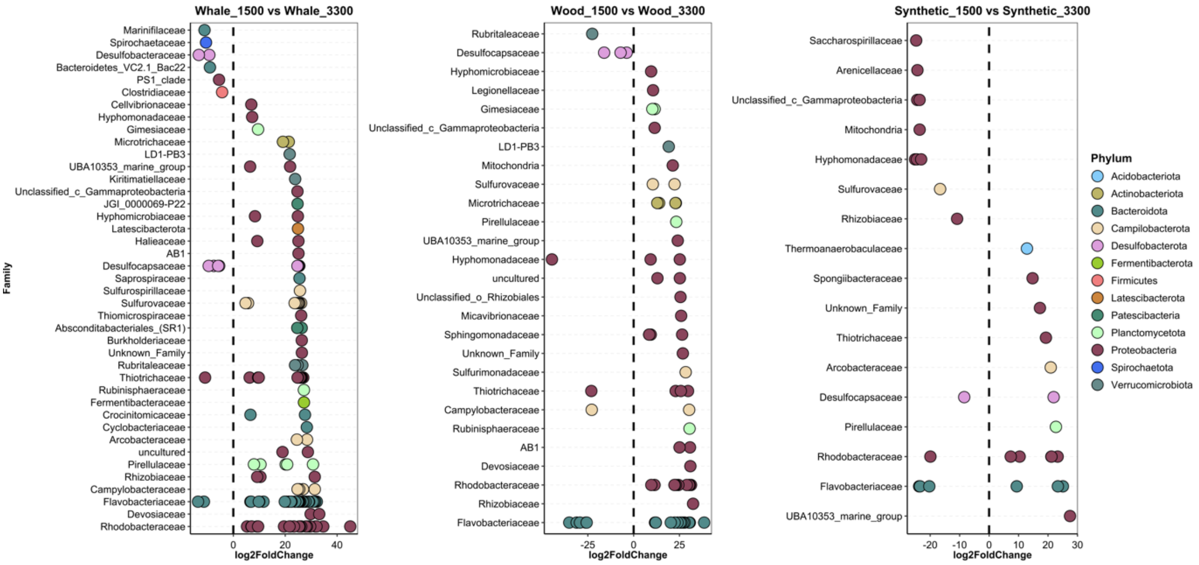


**Figure S4**. Differential abundance analysis of the ASVs between the same substrate type at different sites (1500 and 3300 m isobaths). Log2FoldChange < 0 and > 0 indicates, respectively, ASVs with significantly lower and higher abundance in site 1 vs site 2:  A) whale 1500 vs whale 3300; B) wood 1500 vs wood 3300; C) synthetic 1500 vs synthetic 3300. The ASVs are grouped by family and coloured by phylum (legend).

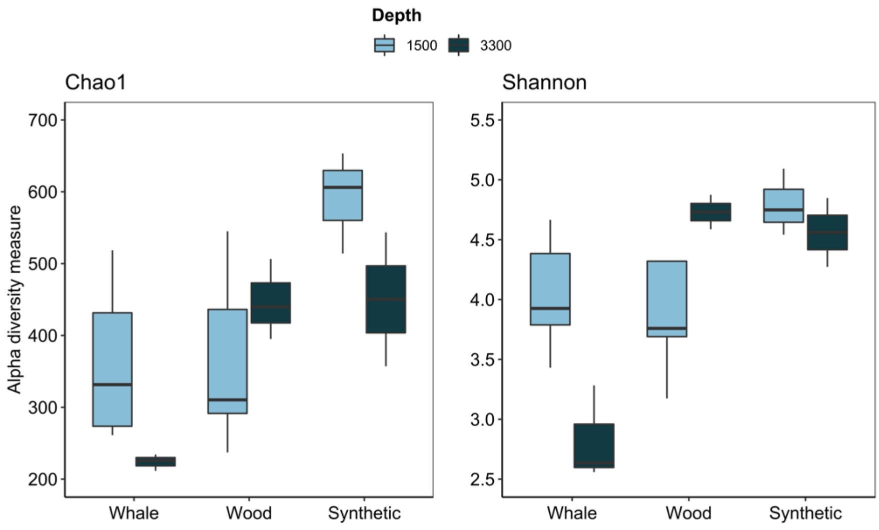


**Figure S5**. Alpha diversity of the prokaryotic communities in the different substrates at the depths 1500 and 3300 meters.

**Table S1**. Environmental characteristics of the sites and number of samples collected

| Isobath | Temperature | DO | Pressure | Salinity | Conductivity | GPS | | Number of samples collected | | |
|---|---|---|---|---|---|---|---|---|---|---|
| | | | | | | Lat | Long | WH | WO | SY |
| 1500 | 3.6 | 3.074 | 1413.5 | 34.7 | 3.264 | -21.45010 | -39.896583 | 7 | 5 | 3 |
| 3300 | 2.5 | 4.012 | 3156.7 | 34.9 | 3.246 | -22.84086 | -38.416333 | 3 | 3 | 2 |

WH - whale vertebrae; WO - wood; SY – synthetic

**Table S2**. Two-way ANOVA statistics evaluating the effects of depth and substrate type on Chao1 and Shannon diversity indexes.

| **Factor** | **Chao1** | | **Shannon** | |
|---|---|---|---|---|
| | F | *p value* | F | *p value* |
| depth | 1.02 | 0.327 | 1.03 | 0.324 |
| substrate | 7.97 | **0.004** | 7.37 | **0.005** |
| depth * substrate | 3.40 | 0.057 | 7.33 | **0.005** |
